# Genomic Insights of an Andean Multi-resistant Soil Actinobacterium of Biotechnological Interest

**DOI:** 10.1101/2020.12.21.423370

**Authors:** Daniel Alonso-Reyes, Fátima Silvina Galván, Luciano Raúl Portero, Natalia Noelia Alvarado, María Eugenia Farías, Martín P. Vazquez, Virginia Helena Albarracín

## Abstract

Central Andean Ecosystems (between 2000 and 6000 masl) are typical arid to semiarid environments suffering from the highest total solar and UVB radiation on the planet but displaying numerous salt flats and shallow lakes. Isolated from these environments, Andean Microbial Communities (AME) of exceptional biodiversity endures multiple severe conditions. Also, the poly-extremophilic nature of AME’s microbes indicates the potential for biotechnological applications. In this context, the presented study used genome mining and physiological characterization to reveal the multi-resistant profile of *Nesterenkonia sp*. Act20, an actinobacterium isolated from the soil surrounding Lake Socompa, Salta, Argentina (3570 m). UV-B, desiccation, and copper assays showed the strain’s exceptional resistance to all these factors. Act20’s genome presented coding sequences involving antibiotics, low temperatures, UV and arsenic resistance, nutrient limiting conditions, osmotic stress response, low atmospheric oxygen pressure, heavy metal stress, and resistance to fluoride and chlorite. Act20 can also synthesize proteins and natural products such as an insecticide, bacterial cellulose, ectoine, bacterial hemoglobin, and even antibiotics like colicin V and aurachin C. We also found numerous enzymes for animal and vegetal biomass degradation and application in other industrial processes.

The herein report shed light on the microbial adaptation to high-altitude environments, its possible extrapolation for studying other extreme environments of relevance, and its application to industrial and biotechnological processes.

**HIGHLIGHTS:**

- Arid Andean Soils are attractive sources of microbial strains useful in biotechnological processes.
- Physiological studies revealed the multi-resistant nature of the poly-extremophile *Nesterenkonia sp*. Act20.
- Act20’s genome analysis showed a complete set of genes coding for proteins involved in resistance to multiple stresses, including extremoenzymes and extremolytes.

## INTRODUCTION

Central Andean Ecosystems (between 2000 and 6000 masl) are typical arid to semiarid environments suffering from the highest total solar and UVB radiation on the planet, displaying numerous salt flats and shallow lakes (Albarracín et al., 2016, 2015). Spanning from the Atacama Desert in Chile, through the Argentinean and Bolivian Puna up to the Peruvian Andes, these ecosystems experience a wide daily temperature range, high salinity (up to 30%), scarce nutrient availability and high concentration of heavy metals and metalloids, especially arsenic (Albarracín et al., 2016, 2015).

Despite these conditions, Andean Microbial Communities (AME) prove exceptional biodiversity and diverse strategies for enduring these severe conditions (Albarracín et al., 2016, 2015; Solon et al., 2018). Likewise, the importance of exploiting AME poly-extremophiles’ full potential in terms of their biotechnological applications was highlighted (Albarraci and Farías, 2012). Examples are the production of waxes and fatty acids for biodiesel (Bequer Urbano et al., 2013) or compatible solutes, antioxidants, pigments, or enzymes for the pharmaceutical industry (Farias et al., 2011). Current projects heading this way have yielded detailed molecular information and functional proof on novel extremoenzymes: i.e., photolyase of *Acinetobacter sp*. Ver3 (Albarracín et al., 2014), an arsenical resistance efflux pump, and a green tuned microbial rhodopsin (Albarracin et al., 2015) in *Exiguobacterium sp*. S17 (Ordoñez et al., 2015).

*Actinobacteria* are high GC (50-71 %), Gram-positive microbes found in both terrestrial and aquatic ecosystems (Albarracín et al., 2005; Montalvo et al., 2005). Being the soil microbiota’s main component (Schrempf, 2013), Actinobacteria exhibit various morphologies (Ventura et al., 2007), physiological and metabolic properties, and includes many species, which are useful in biotechnology (Kurtböke, 2003). Previous work using dependent and independent culture techniques revealed that *Actinobacteria* is one of the predominant taxonomical groups among the AME’s microbial communities (Dib et al., 2008, 2009; Rasuk et al., 2017). Moreover, AME’s *Actinobacteria* have been demonstrated to carry giant linear plasmids that may involve the community’s spread of resistance traits (Dib et al., 2010). AME’s actinobacteria also showed their potential for producing secondary metabolites useful for the pharmacy industry. Wichner et al. (2017) reported that the extremotolerant isolate, *Lentzea* sp H45, synthesized new monoene, and diene glycosides. These natural products called lentzeosides A-F possess inhibitory activity against HIV integrase, a key enzyme for recombining the HIV genome into the host genome. Schulz et al. (2011) evidenced the production of bioactive compounds called abenquines (simple aminobenzoquinones substituted by different amino acids) by *Streptomyces* sp. DB634. Abenquines showed moderate inhibitory activity against phosphodiesterase type 4 (PDE4b) while proved useful for treating inflammatory diseases. Moreover, *Streptomyces* C38 produces 22-membered macrolactonic antibiotics atacamycin A-C, also considered drugs for treating inflammatory diseases by inhibiting PDE4b and antitumor by acting against tumor cell lines (Nachtigall et al., 2011).

*Nesterenkonia* is a particular genus (Stackebrandt et al., 1995) with most representatives isolated from hypersaline or alkaline environments such as saline soil, solar salt, seafood, soda lake, or alkaline wastewater. Halophiles are exploitable microorganisms for bioprocesses (Fu et al., 2014; Liu et al., 2019; Yin et al., 2014) (Yue et al., 2014). For example, *Nesterenkonia MSA 31* isolated from a marine sponge *Fasciospongia cavernosa* produces a halo-alkali and thermal tolerant biosurfactant useful as an emulsifier stabilizing agent in the food industry (Kiran et al., 2017). Furthermore, *Nesterenkonia* sp. strain F isolated from Aran-Bidgol Lake (Iran) can produce acetone, butanol, ethanol, acetic, and butyric acids under aerobic and anaerobic conditions (Amiri et al., 2016). Also, *Nesterenkonia lacusekhoensis EMLA3* degrades reactive violet 1 (RV1), a toxic azo dye, under conditions of high pH and in the presence of a high concentration of NaCl, both of which generally inhibit microbial treatment process (Prabhakar et al., 2019). Thus, *Nesterenkonia* strains are attractive microbes in the search for biotechnological resources.

In a previous screening for extremophilic *Actinobacteria* from Puna arid alkaline soil, the dark yellow-pigmented *Nesterenkonia sp*. Act20 strain was identified (Rasuk et al., 2017). Act20 grew at a high concentration of NaCl (25%), Na_2_CO_3_ (5mM), and arsenic (up to 200mM arsenate), in a wide range of pH 5-12, with optimal growth at alkaline pH (Rasuk et al., 2017). In subsequent works, our group evidenced that *Nesterenkonia* sp Act20 had a high tolerance to UV-B (up to 100 kJ/m^2^) due to an integrated response to radiation called UV-resistome (Portero et al., 2019).

The following work aims to test the multi-resistance of the Act20 strain combining phenotypic profiling with in-depth genomic analysis. Also, we highlight the potential for biotechnological use of extremozymes and extremolytes coded in its genome.

## 1. MATERIALS AND METHODS

### 2.1. Strains and culture conditions

UV-resistant strain Act20 used in this study was previously isolated from soil around Lake Socompa (3,570 m) at the Andean Puna in Argentina (Albarracín et al., 2016, 2015) and belongs to the LIMLA-PROIMI Extremophilic Strain Collection. Bacterial strain *Nesterenkonia halotolerans* DSM 15474 belongs to DSMZ Bacterial Culture Collection, and we used it as a control following previously reported works (Rasuk et al., 2017). Both strains were grown in an “H” medium (a medium modified for halophiles, containing NaCl 15 g L^-1^, KCl 3 g L^-1^, MgSO_4_ 5 g L^-1^, sodium citrate 3 g L^-1^) added with 2% agar when applicable.

### 2.2. Multi-resistance assays

The resistance of strain Act20 cells to diverse physical and chemical stresses was tested when exposed to increasing concentrations of copper, high UV doses, and desiccation treatments. For studying the response to desiccation, cells were first grown aerobically at 30 °C overnight in a 10 ml nutrient broth medium on a rotary shaker. Cells were harvested, washed once with sterile NaCl solution (0.85 %, w/v), and resuspended to reach an OD_600nm_ of 2 (±0.1) in NaCl solution. Approximately 20 aliquots (100 μl each) from this cell suspension were spotted onto 0.45-μm filters (Sartorius, Göttingen, Germany). These filters were placed onto agar medium H plates and incubated at 30 °C for five days. The colonies were then let dry by incubation in empty sterile dishes at 25 °C and 18% of relative humidity for 50 days, and the viable count (CFU) was assessed at different times. The filters on which the strains were grown were added to sterile microcentrifuge tubes. The cells from each filter were resuspended separately, and the CFU was determined before and after desiccation treatment. The tolerance to desiccation was determined in Act20 and *N. halotolerans* DSM 15474 as control.

Resistance to UV irradiation and copper salts were tested by a quick qualitative method. For the copper resistance profile assays, aliquots of 5 μL of an overnight (OD_600_≈0.6) culture were loaded onto H medium agar plates supplemented with 1, 2, or 3 mM CuSO_4_. The control cultures consisted of an “H” medium without copper supplementation. Then they were incubated for 72 h at 30 °C under continuous PAR luminosity conditions with an OSRAM 100 W lamp.

For the UV resistance assays, the cells were pre-culture on liquid medium H, and once at OD_600nm_ = 0,6 collected for serial dilutions. Aliquots of 5 μL were then loaded onto medium agar plates and immediately exposed to UV-B irradiation (Vilbert Lourmat VL-4, the maximum intensity at 312 nm) 5 min (1,7 Kj m^-2^), 15 min (5,1 Kj m^-2^), and 30 min (10,4 Kj m^-2^). Then they were incubated for 72 h at 30 °C in the dark to prevent photoreactivation. UV-B irradiance was quantified with a radiometer (Vilbert Lourmat model VLX-3W) coupled with a UV-B sensor (Vilbert Lourmat model CX-312). The minimal intensity measured was 5,21 W m^-2^, and maximal power was 5,4 W m^-2^. Controls of unexposed samples were run simultaneously in the dark.

Microbial growth was recorded with tree signs (+++) when similar to the growth in controls, two signs (++) when it was slightly different from the growth in the controls, one sign (1 pts) when growth was low (isolated colonies), and no sign when it was no growth at all. Parallel assays were performed for *Nesterenkonia halotolerans* DSMZ 15474 to compare resistance profiles.

### 2.3. Microscopic observation and ultrastructural characterization of Act20 cells

These assays were designed to evaluate the morphology and ultrastructure of Act20 in challenging conditions similar to those present in their original environment, i.e., under high UV irradiation and chemical (copper) stress. The selected strains were grown in H medium at 30°C with shaking (180 rpm). Cells in the mid-log phase of growth were harvested by centrifugation (5000 rpm for 10 min). Pellets were washed twice in 0.9 % NaCl and were kept under starvation conditions for 18 h at 4 °C in the same solution. 20 ml of cell suspension was transferred to a sterile plate and were exposed to UV-B irradiation at different times, as indicated before. Copper-challenged cultures were likewise obtained by growing the cells in H medium with and without 3 mM Cu.

For scanning electron microscopy (SEM) and transmission electron microscopy (TEM), 100 μL aliquots were collected for each different treatment and centrifuged (5,000 rpm for 10 min) to remove the supernatant. The pellets were immediately fixed with Karnovsky’s fixative (a mixture of 2,66% paraformaldehyde and 1,66% glutaraldehyde) in a 0.1 M phosphate buffer pH 7.3, for 48 h at 4°C. For SEM, the cells were processed according to previously optimized methods (Zannier et al., 2019). Briefly, aliquots of 50 μl of samples fixed were placed in coverslips for electron microscopy and kept for three hours at room temperature. The samples were then dehydrated in graded ethanol (30%, 50%, 70%, 90%, and 100%) for 10 min each and finally maintained in acetone 100% for 40 min. The dehydration was completed with the critical drying point (Denton Vacuum model DCP-1), in which acetone was exchanged by liquid CO_2_. Then, samples were mounted on stubs and covered by gold (Ion Sputter Marca JEOL model JFC-1100) and observed under a Zeiss Supra 55VP (Carl Zeiss NTS GmbH, Germany) scanning electron microscope belonging to the Electron Microscopy Core Facility (CIME). For TEM, the protocol from Albarracín et al. (2008) was followed. After fixation, samples were washed twice in 0.1 M phosphate buffer, pH 7.3 (5000 rpm for 10 min), and embedded in agar (Bozzola, 2007). Agar pellets were post-fixed in 1% osmium tetroxide in phosphate buffer, pH 7.3, overnight at 4°C. After washing with the same buffer, and were stained in 2% uranyl acetate solution for 30 min at room temperature. The samples were dehydrated with ethanol solutions increasing concentrations (70%, 90%, and 100%) for 15 min each and finally maintained in acetone 100% for 30 min. After that, the infiltration and embedding in an acetone-SPURR resin sequence were carried out, followed by polymerization at 60°C for 24 h. Ultrathin sections were cut using a diamond knife on a manual ultramicrotome (Sorvall Porter-Blum Ultramicrotome MT-1). Bacteria were examined using a Zeiss LIBRA 120 (Carl Zeiss AG, Alemania) transmission electron microscope at 80 kV, belonging to the Electron Microscopy Core Facility (CIME-CONICET-UNT).

### 2.4. Genome sequencing, assembly, and gap closure

Genomic DNA from *Nesterenkonia sp*. Act20 strain was purified from cells grown on LB broth for 72 h at 30° C and harvested by centrifugation (3,000 g for 10 min at 4°C). Pellets were washed twice with distilled water. We extracted total genomic DNA with the DNeasy Blood and Tissue Kit (Qiagen) following the manufacturer’s recommendations. Whole-genome shotgun pyrosequencing was achieved using a 454 preparation kit (Roche Applied Sciences, Indianapolis, IN, USA) and sequenced with a GS-FLX using Titanium chemistry (454 Life Sciences, Roche Applied Sciences). The 454 reads were assembled with Newbler Assembler software, v. 2.5.3, with-URT option. Extra-assembling programs were run: MIRA v. 3.4.0 and Celera Assembler, v. 6.1. The different assemblages were fused using MINIMUS 2 Pipeline from the AMOS Package. The merged assembly was used as a guide for designing the primers, which were, in turn, used to confirm contig joints and close gaps. The overall sequence coverage was 37X; This Whole Genome Shotgun project has been deposited at DDBJ/ENA/GenBank under the accession JADPQH000000000. The version described in this paper is version JADPQH010000000.

### 2.5. Genome analyses

Genome annotation was implemented using PROKKA (Seemann, 2014) with a custom expanded protein database, which includes Swiss-Prot, TrEMBL, Pfam, SUPERFAMILY, TIGRFAM, and a genus-specific database built from GenBank files downloaded from the NCBI database. We processed the set of annotated FASTA files belonging to genomes with Proteinortho, which detects orthologous genes within different species. For doing so, the software compares similarities of given gene sequences and clusters them to find significant groups (Lechner et al., 2011).

### 2.6. Phylogenetic analysis

The sequence-based taxonomic analysis was performed using both 16S rDNA and whole-genome comparisons. Sequences from genomes and 16S rDNA were obtained from the NCBI database (Assembly and RefSeq), and their characteristics resumed in Supplementary Table S1. Some 16S rDNA were retrieved from genome sequences to link both types of analysis. Sequences from 16S rDNA were aligned with Silva Incremental Aligner (Pruesse et al., 2012), and the phylogenetic tree was created with Fasttree 2.1.7 (Price et al., 2010) with the Maximum Likelihood method using Generalized Time-Reversible (GTR) model. Subsequent processing and visualization of the tree were performed with iTOL (Letunic and Bork, 2016). Genome-wide nucleotide identity trends were explored in the genome dataset by estimating all-against-all pairwise Average Nucleotide Identity (ANI). We utilized the ANIm approach that uses MUMmer (NUCmer) to align the input sequences as implemented in pyANI (Kurtz et al., 2004). The average between any given pair was used as the final value. Heat maps were generated using the heatmap V 1.0.12 R package (Kolde, 2019).

## 2. RESULTS

### 3.1. Taxonomic Affiliation of *Nesterenkonia* sp. Act20

*Nesterenkonia sp*. Act 20 was first isolated by Rasuk et al. (2017) and initially assigned to the genus by partial 16S rDNA sequencing. This work compared the full 16S rDNA sequence from Act20 with related strains, resulting in 98.2 % identity with *N. sandarakina* YIM 70009 and 97.4 % identity with both *N. jeotgali* JG-241 and N. *halotolerans* YIM70084. The phylogenetic tree based on this marker also suggests a close relationship of Act20 with the above mentioned plus *N. sp*. AN1, *N. aurantiaca* strains, *N. sandarakina,* and *N. Lutea*, clustering together in a significant clade (Fig. 1A). The whole-genome analysis using the ANI method shows a similar pattern of relationship (Fig. 1B). In this analysis, *N. sp*. Act 20, *N. aurantiaca* DSM 27373, *N. sp*. AN1, *N. jeotgali* CD087, and *N. sandarakina* CG 35 cluster with significance, and Act20 have a lower percent average nucleotide difference (defined as 100% - ANI) value than the proposed 95% threshold (Richter and Rosselló-Móra, 2009), suggesting that it could be a novel species.

**Figure 1.**
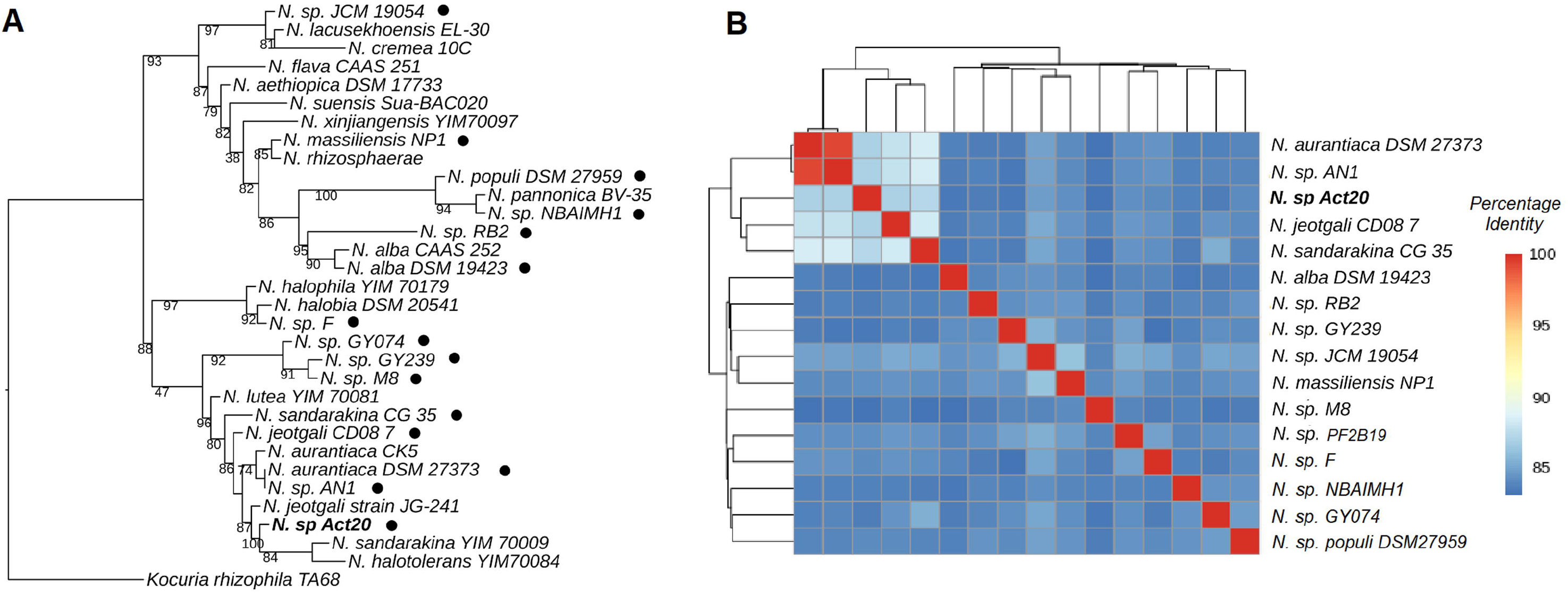
Phylogenetic analysis of *Nesterenkonia* strains. Strain *Nesterenkonia* sp. Act20 is in bold. (A) Maximum likelihood tree from the 16S rRNA gene. Black dots indicate that the gene was retrieved from a sequenced genome. (B) Heatmap based on whole-genome average nucleotide identity (ANI). *Act20* is distinct and inside the significant cluster.

### 3.2. Multi-Resistance Profile of Act20

Resistance to desiccation, UV, and copper was uncovered in *Nesterenkonia* sp. Act20 and compared with the closest relative, *N. halotolerans* (NH). A summary of this multi-resistance is presented in Table 1. Tolerance to desiccation was tested every seven days for seven weeks. Act20 maintained its population in the same order of magnitude for 14 days, but from the third week on, it showed null development (Fig. S1). In turn, NH decreased its population in one order of magnitude after 7 and 14 days of treatment while it did not survive beyond 14 days of continuous drying conditions.

**Table 1.** Multi resistance profile of Act20 strain and comparison with Nesterenkonia halotolerans (NH)

|  | Act20 | NH |
| --- | --- | --- |
| Desiccation Period |  |  |
| **7-days** | +++ | + |
| **14-days** | ++ | + |
| **21-days** | - | - |

| **Copper Concentration** |  |  |
| --- | --- | --- |
| **1 mM** | +++ | +++ |
| **2 mM** | ++ | ++ |
| **3 mM** | + | + |

| **UV Dose** |  |  |
| --- | --- | --- |
| **1,7 Kj m<sup>-2</sup>** | +++ | ++ |
| **5.1 Kj m<sup>-2</sup>** | ++ | + |
| **10.4 Kj m<sup>-2</sup>** | + | + |

Tolerance to UV radiation was tested by placing culture serial dilutions drops of the studied strains on “H” agar plates and exposing them to UV source as described previously. A similar procedure was tried for testing resistance profiles to copper salts in media amended with 1, 2, and 3 mM of copper. Our results showed that Act20 was much more resistant to radiation than the selected control strain (Fig. S2), growing even after a dose of 10,4 Kj m^-2^ of UV-B radiation. In turn, the copper resistance profile was similarly high for both strains; Act20 and NH developed quite well even at the maximum concentration tested (3 mM).

### 3.3. General Genomic Features

The assembly process led to two scaffolds: the largest with 2,092,188 bp in length and G + C content of 65.98%, and the second scaffold with 836,993 bp and G + C content of 65.82%. As a whole, the genome of *Nesterenkonia* sp Act20 consists of 2,930,097 bp, with a GC content of 65.9%. PROKKA annotation shows 2,672 coding sequences, including 2,377 annotated genes and 58 RNAs. Act20 genomics features were compared to the other fifteen *Nesterenkonia* genomes available in the NCBI assembly database (Table S7). Supplementary Table S2 shows the annotated genes of Act20, their sequence length, and functions assigned by homology, as well as the KO identifiers linked for some of these genes. A summary of the most relevant annotated functions can be found in Supplementary Tables S3 and S4.

### 3.4. Genome traits of Act20 multi-resistance phenotype

Genome inspection indicated genetic determinants coding for systems potentially involved in the high resistance profile observed in herein described and previous lab assays (Rasuk et al., 2017). The genome showed pathways associated with osmotic and oxidative stress response, low temperature, starvation response, and low oxygen conditions, unmistakable evidence of this microbe adaptation to its extreme and changing habitat. It also has traits related to resistance to heavy metals (mainly copper and mercury), antibiotics (mainly beta-lactams and vancomycin), arsenic, fluoride, and chlorite (Table S3, S4).

Both, the Prokka and RAST annotations account for several genes for resistance/tolerance to copper (Table 2). Among the direct mechanisms of resilience to copper described by Giachino and Waldron (2020) in Act20 is worth to mention the *cop* family, implicated in copper homeostasis through the capture and expulsion of Cu[I] ions from the cytosol to the periplasm. This mechanism may be complemented by the action of a copper oxidase necessary for converting Cu [I] to Cu [II], which is more biologically inert and tend to remain in the periplasm. It is also interesting the presence of a gene matching a recently discovered family of copper-binding proteins involved in cytosolic copper storage (Dennison et al., 2018). On the other hand, the indirect mechanisms would involve the participation of genes whose actions compensate for specific damage caused by toxic copper. This is the case for the specific DsbD oxidase which rearranges misfolded peptides, and the Fur master regulator of iron metabolism that counteracts the constant scavenging of iron cofactors from enzymes, which in turn can generate ROS (Giachino and Waldron, 2020).

**Table 2.** Group of genes potentially involved in the copper resistance reported for Act20.

| **Copper tolerance/resistance trait** | **Function** |
| --- | --- |
| **Copper-translocating P-type ATPase** | Probably similar to P-type ATPase CopA, as secretes Cu(I) from the cytoplasm to the homeostasis (Giachino and Waldron, 2020). |
| **CopC domain protein** | CopC is a bacterial blue copper protein that molecule. Along with CopA, CopC mediates copper in the periplasm (Arnesano et al., 2003) |
| **Putative CopD protein** | Exact function not known. Involved in copper uptake in conjunction with CopC (Swiss) |
| **CopZ** | Chaperone that serves for the intracellular sequence from the cytosol to the periplasm (Giachino and |
| **Copper oxidase** | Probably a periplasmic copper oxidase-like C the cytosol by converting periplasmic Cu(I) to inert, making it a relatively “safe” species for c (Waldron, 2020). |
| **DsbD domain-containing protein** | DsbD (also known as CutA) oxidoreductase re incorrect disulfide bonds are inserted by toxic c |
| **Four-helix bundle copper-binding protein** | Cu(I) cytosolic storage. First found in meth amounts of copper to metabolize methane. Its what they are storing copper for remains unknow |
| **Fur family ferric uptake transcriptional regulator** | As toxic copper provokes mismetalation o regulator of iron metabolism is overexpressed i copper resistance even at low copper concentra |

Previous assays carried out on Act20 revealed its ability to grow in alkaline media (up to pH 12) and at high NaCl (25%) and Na_2_CO_3_ (5 mM) concentrations (Rasuk et al., 2017). In this work, we also verify its desiccation tolerance, a phenotype that is supported by a vast repertory of genes (Table 3). This ability may be explained by the presence of transporters for the uptake of a diverse organic osmoprotectant such as glycine betaine, proline betaine, glycerol, choline, and trehalose. Genes for the synthesis of glycine betaine and the complete set of genes for ectoine were also detected (Table 3, Fig. S3). These compounds counteract the environment’s high osmolality avoiding a rapid efflux of water from the cell and, consequently, the loss of turgor. Low turgor starts a rapid influx/synthesis of these osmo-protectants that complement another inorganic compounds (Lucht and Bremer, 1994; Nagata and Wang, 2001; Styrvold and Strom, 1991; Wood, 1988). Other genes, like OsmC and MdoB, are involved in coping the osmotic stress in peculiar ways (Atichartpongkul et al., 2001), specifically MdoB, whose product requires periplasm and outer membrane facilitate its effect (Sleator and Hill, 2002), both structures not normally present in Gram-positive bacteria including Act20.

**Table 3.**
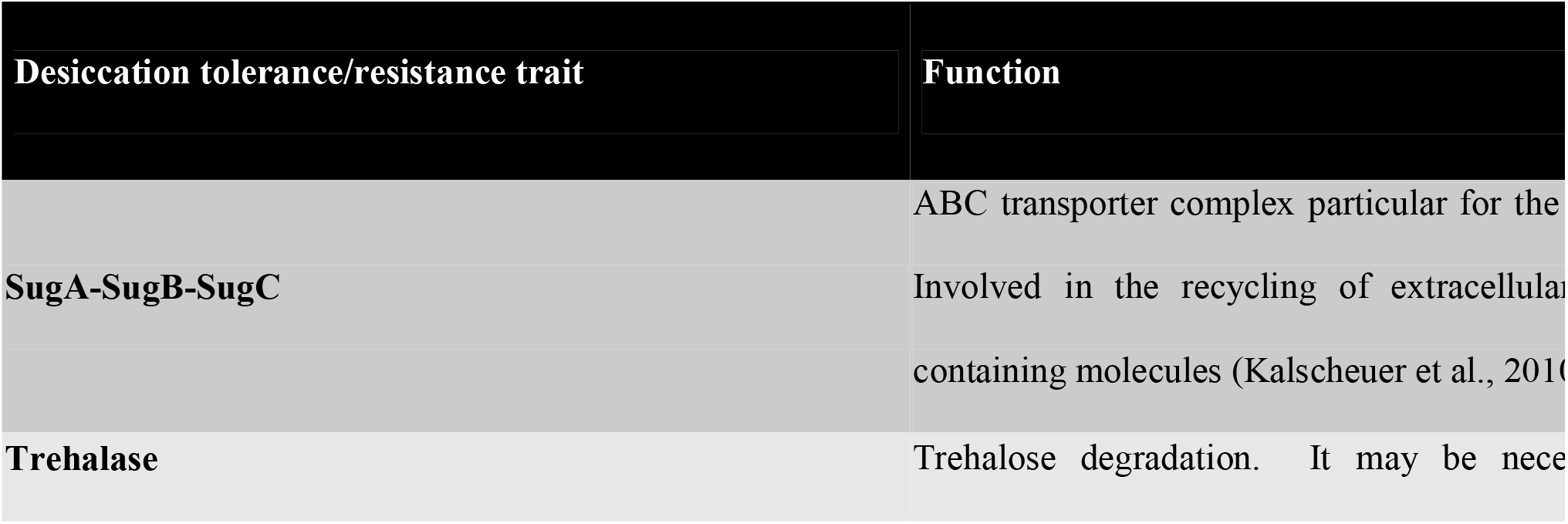
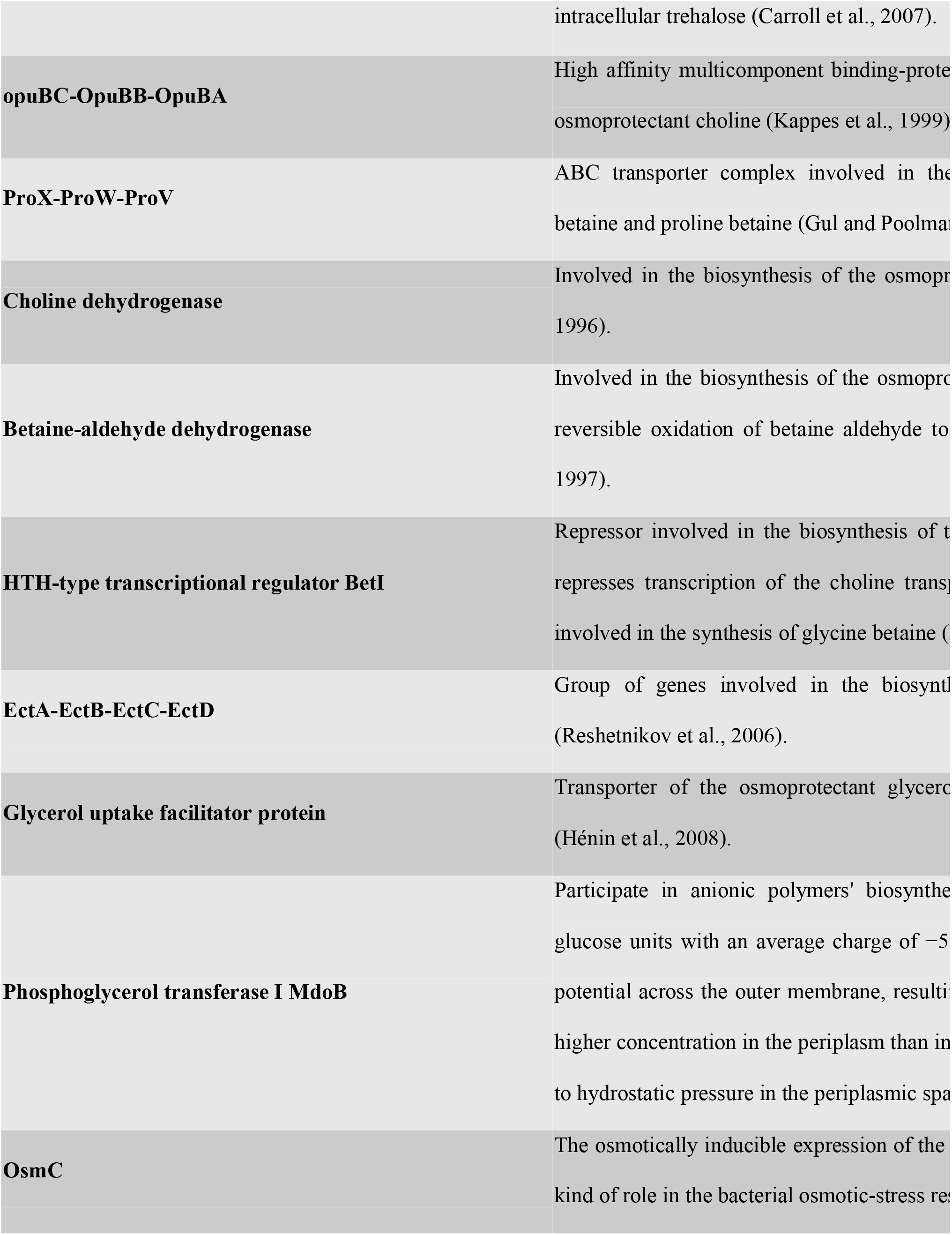
Group of genes potentially involved in the desiccation resistance reported for Act20.

The genomic profile of ABC transporters and the two-component system is also a reflection of the original environment. The set of proteins to sense phosphate limitation and subsequent phosphorus incorporation (in the form of phosphates and phosphonates) are evident (Fig. S4, S5). A similar is noticed for osmotic stress. Other environmental sensing types include oxygen limitation, low temperature, cell envelope stress, cell wall stress, and antibiotics. Transporters for iron and various organic compounds such as carbohydrates (mostly of plant origin), nucleosides, amino acids, and oligopeptides were also reported.

To discover unique functional traits of Act20, we searched the orthologous sequences for public available *Nesterenkonia* genomes. The set of annotated FASTA files belonging to genomes were processed through Proteinortho, which detect orthologous genes within different species. The proteins of Act20 that did not match any ortholog were analyzed one by one to verify if the annotated function they encode is unique for Act20 and is not present in other representatives of the genera. Table 4 shows the exclusive traits of Act20, most of them related to its extreme natural environments such as bacterial persistence, bacterial cell envelope stress response, and resistance to osmotic stress, desiccation, and phosphate starvation. Degradative enzymes 6-deoxy-6-sulfogluconolactonase and α-xylosidase take part in the decomposition of prototrophic biomass present in the soil, from which Act20 could take advantage. The remaining functions were characterized only at the protein domain level, like ethyl tert-butyl ether degradation, second messenger’s sensors, and cell wall binding.

**Table 4.** New functional traits (with no counterpart in other *Nesterenkonia*) of *Nesterenkonia sp. Act 20* unveiled by genome functional annotation and ortholog clustering.

| Unique trait | Function |
| --- | --- |
| **Protein SphX** | High-affinity phosphate-binding protein induced by phosphate starvation (Gobler and Gobler, 2013) |
| **Serine/threonine-protein kinase toxin HipA** | Toxin that induces bacterial persistence and is essential for survival in stationary phase (Gobler, 2013). |
| **6-deoxy-6-sulfogluconolactonase** | Sulfoquinovose degradation (Felux et al., 2015) |
| **Trehalase** | Trehalose degradation (Carroll et al., 2007). |
| **α-xylosidase** | Catalyzes the liberation of alpha-xylose from the xyloglucan oligosaccharides (Matsuzawa et al., 2007). |
| **4-deoxy-L-threo-5-hexosulose-uronate ketol-isomerase** | Plays a role in the catabolism of hexuronate and is likely substituting for the regular hexuronate isomerase. Expression of this gene is repressed in these conditions (Rochet et al., 2010). |
| **Pleckstrin homology domain (bPH_1)** | Appears to be involved in the bacterial cell cycle (Rochet et al., 2010). |
| **SMODS-associated and fused to various effectors sensor (SAVED) domain** | Senses nucleotide or nucleotide derivatives and is involved in the regulation of effectors deployed by a class of conflict systems. It is involved in the sensing of the nucleotide second messengers (Fischer et al., 2010). |
| **WxL domain** | Cell wall binding in gram-positive bacteria. It is involved in the binding to peptidoglycan (Brinster et al., 2007). |
| **EthD domain** | Related to the EthD protein, which is involved in the regulation of ether (ETBE) (Chauvaux et al., 2001) |
| **Coronavirus endopeptidase C30** | Involved in viral polyprotein processing in replication (Fischer et al., 2010) |

### 3.5. UV-Resistome of Act20

The high UV-resistance profile of Act20 calls for more in-depth characterization and points out the existence of integrated physiological and molecular mechanisms triggered by ultraviolet light exposure. We named this system “UV-resistome” as described before for other AME poly-extremophiles (Kurth et al., 2015; Portero et al., 2019). Ideally, the UV-resistome depends on expressing a diverse set of genes devoted to evading or repairing the damage provoked directly or indirectly. Ideally, it encompasses the following subsystems: (1) UV sensing and effective response regulators; (2) avoidance and shielding strategies; (3) damage tolerance and oxidative stress response; and (4) DNA damage repair. Therefore, we screened genes associated with each of these UV-B resistome subsystems for all available genomes of the *Nesterenkonia* genus. This approach unveiled the relative genomic potential of Act20 to defend itself from UV-B radiation, a strain that naturally endures the highest irradiation on the planet. Unlike previous works, we studied the UV-resistance integrally and included genes that could have the potential to generate a UV evasion response or could lessen the negative impact the light, such as motility genes, pilus, and gas vesicles (Damerval et al., 1991).

Table S5 details the UV-resistome for every strain and clusters it by subsystems. The collections summarized all types of damage repair (base excision repair, nucleotide excision repair, mismatch repair, homologous repair, direct repair, homologous repair, direct repair, translesion DNA synthesis factors, and SOS response factors) explicitly, oxidative stress response and UV avoidance/protection mechanisms (synthesis of photoprotective pigments, and genes for flagellum, pilus, gas vesicles, and swarming motility; Fig. 2). The bars are sorted from top to bottom, taking into account the number of different genes for each subsystem. Act20 is positioned at the top with the most diverse and complete UV-resistome, with 114 genes, followed by NBAIMH1 with 107 genes, which also belongs to an extreme altitude environment. Interestingly, other strains from harsh environments with expected high solar radiation, such as AN1 and M8 with 85 and 83 genes, respectively, also have robust UV-resistomes. On the other hand, strains belonging to environments with little or null exposure to solar radiation present less diverse and fewer genes: i.e., *N. alba*. DSM19423, GY074, and RB2 with 56, 63, and 65 genes, respectively.

**Figure 2.**
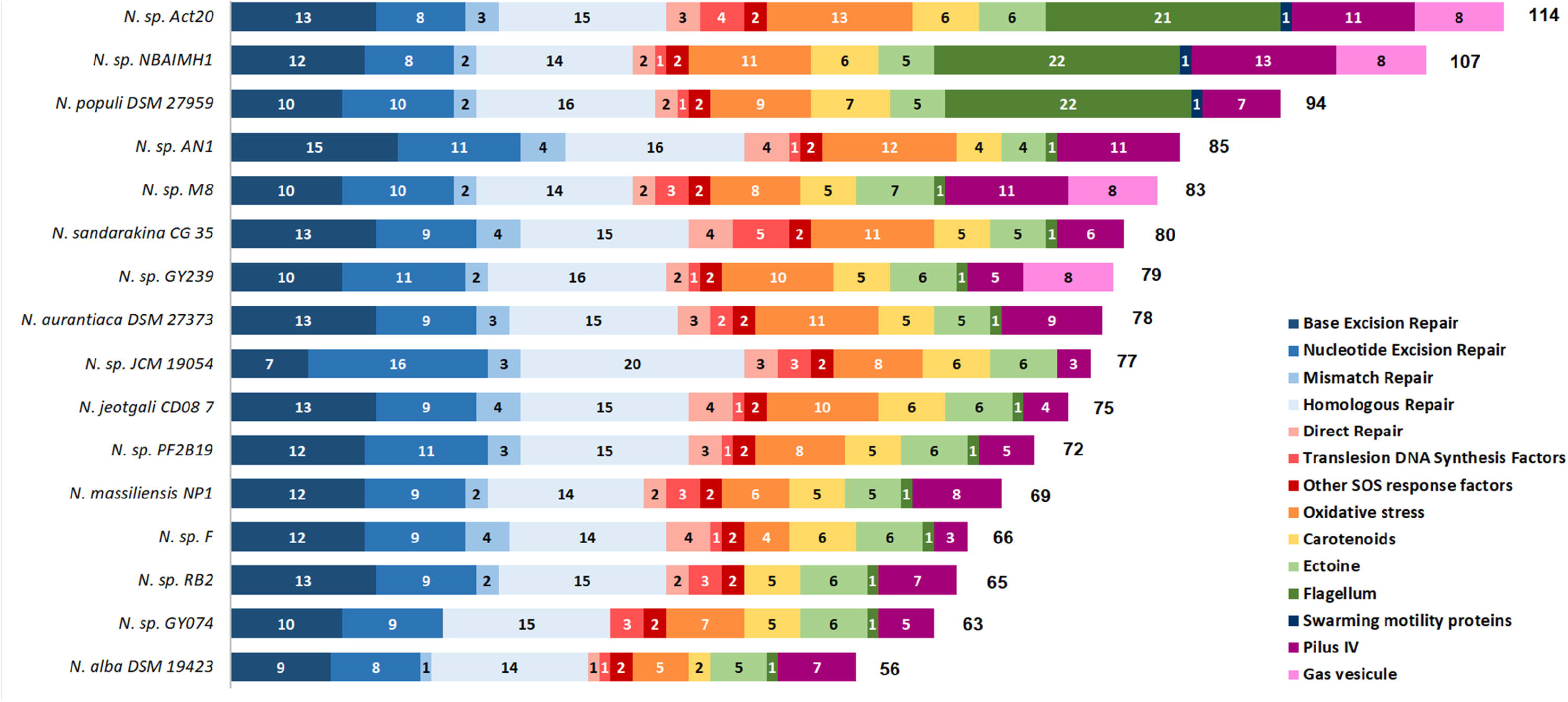
The stacked bar chart shows the count of genes for all UV-resistome subcategories in colors, including the total count at the right end.

### 3.5. Ultrastructural changes in UV-challenged Act20 cells

The morphology and ultrastructure of the actinobacterium *Nesterenkonia* sp. Act20 was observed under UV challenging and non-challenging conditions to determine their behavior in their original environment. *Nesterenkonia halotolerans* DSM 15474 (NH) was used for comparison. Under SEM, in normal conditions, Act20 and DSM 15474 cells appear as irregular coccoids or short rod-shaped. Their size varied between 0.41-0.43 x 0.6-0.82 μm for Act20 and 0.41-0.38 x 0.71-0.51 for NH. The surface was smooth without evidence of cell wall rupture (Fig. 3A, 3E). In contrast, after exposure to UV both strains exhibited morphological alterations, which were consistently more severe in NH cells.

**Figure 3.**
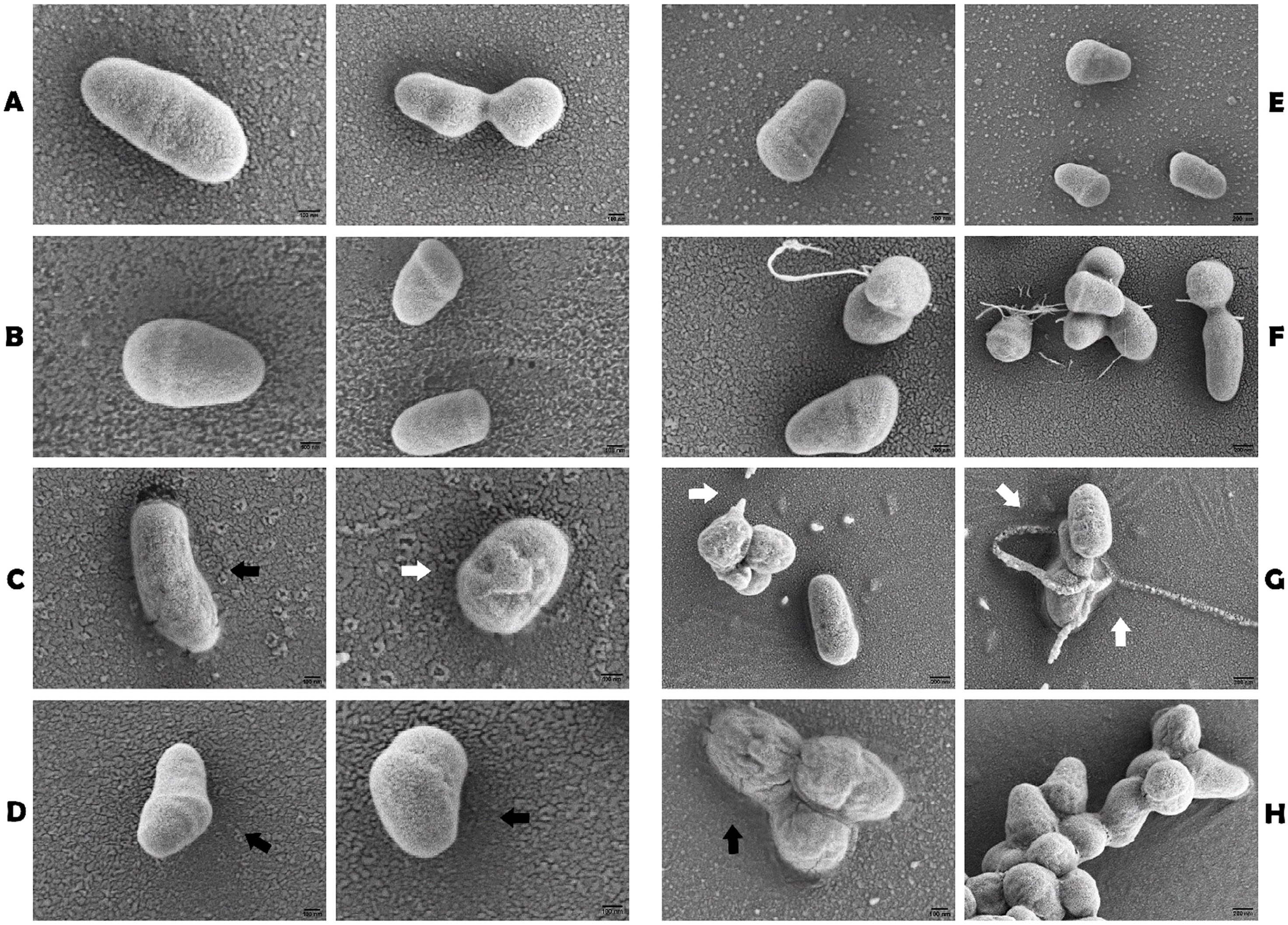
Scanning electron microscopy (SEM) micrographs of actinobacteria after exposure to UV-B radiation. **(A-D)** *Nesterenkonia* sp. Act20. **(E-H)** *Nesterenkonia halotolerans* DSM 15474. **(A, E)** Non-exposed growing bacterial cells (control). **(B, F)** Growing bacterial cells exposed to 0,17 Jls/cm2 of UV-B (5 min). **(C, G)** Growing bacterial cells exposed to 0,51 Jls/cm2 of UV-B (15 min). **(D, H)** Growing bacterial cells exposed to 1,04 Jls/cm2 of UV-B (30 min). Black arrows indicate changes morphological, and white arrows indicate damage in cells **(C)** and pili deterioration **(G)**. Scale bar 200 nm (E, F, G, H), 100 nm (A, B, C, D, E, F, H).

As the dose increases, Act20 morphology changed as individuals became longer and sometimes wide, probably due to cell division’s interruption, causing a complete deformation in the cell (see black arrows in Fig. 3C, 3D). Furthermore, the cells’ surface appeared with shrinkage signs (see white arrows in Fig. 3C). In NH, fibrilar structures were observed only on the surface of cells treated for 5 and 15 min with RUV (Fig. 3F, 3G). We found that at the dose of 0,51 Jls/cm2 (15 min), the pili thickened, and in some sections, they broke or disintegrated (see Fig. 3G, white arrows). At 30 min of exposure (Fig. 3H), cell aggregates were observed in which bacterial cells adhered to one another by self-produced extracellular polymeric (EPS) substances. The surface irregularity was also found, indicating the wall cell rupture and degradation, probably causing cell lysis (Fig. 3H, black arrow).

TEM images for both strains were likewise obtained (Fig. 4). EM analysis revealed the typical Gram-positive bacteria structure, an intense electron opaque inner layer corresponding to the cytoplasmic membrane (CM) and a less electron opaque outer layer or cell wall (CW).

**Figure 4.**
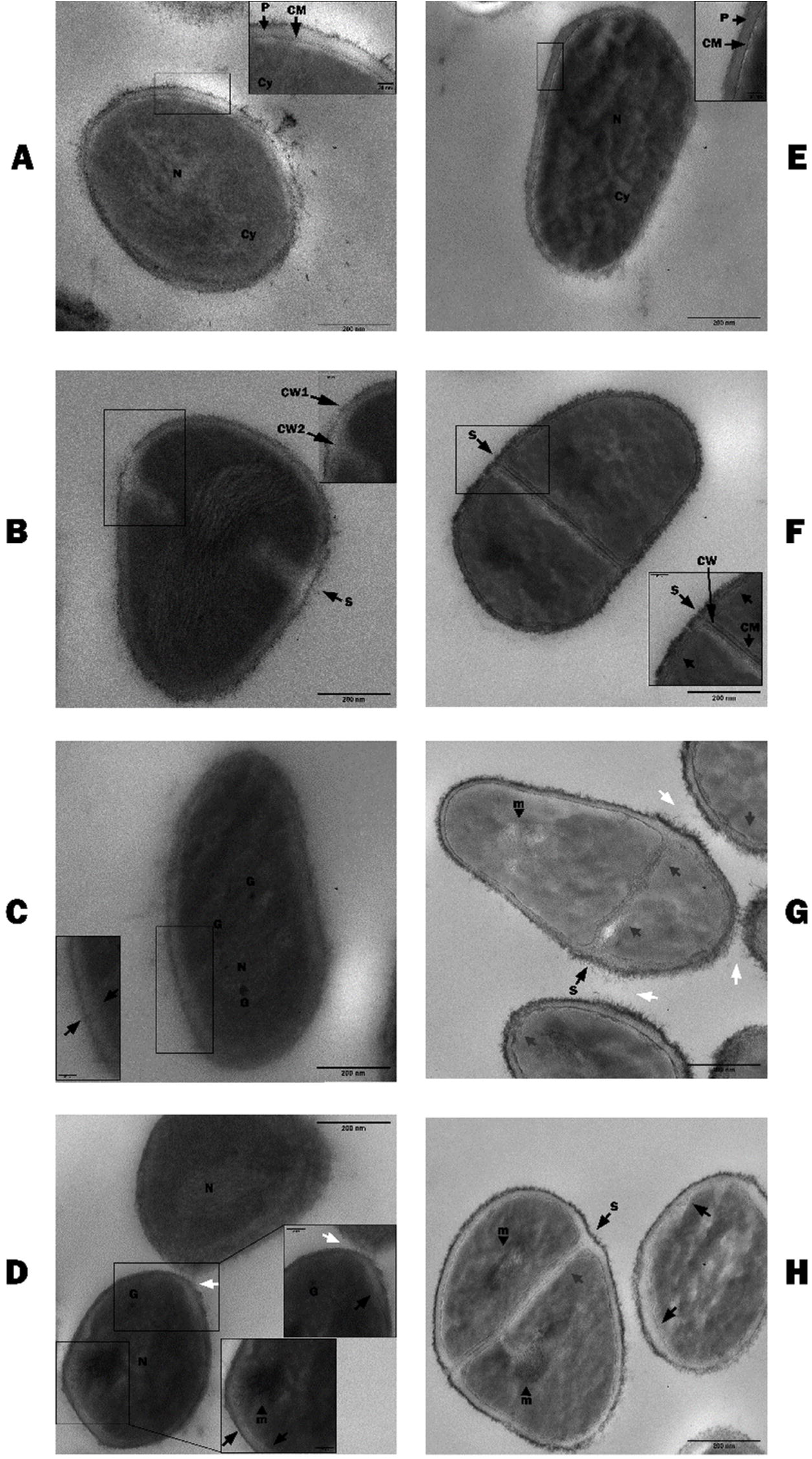
Transmission electron microscopy (TEM) micrographs of actinobacteria cells after exposure to UV-B radiation. **(A-D)** *Nesterenkonia sp*. Act20. **(E-H)** *Nesterenkonia halotolerans* DSM 15474. **(A, E)** Non-exposed growing bacterial cells (control). **(B, F)** Growing bacterial cells exposed to 0,17 Jls/cm2 of UV-B (5 min). **(C, G)** Growing bacterial cells exposed to 0,51 Jls/cm2 of UV-B (15 min). **(D, H)** Growing bacterial cells exposed to 1,04 Jls/cm2 of UV-B (30 min). Scale bar 200 nm principal figure and 50 nm small box.

Ultrastructural changes were observed in the cells after exposure to UV radiation, especially at the structural membrane level, acquiring a certain degree of disorganization compared to the untreated strains (controls). A general observation in both UV-treated bacteria was that numerous cells contained septa (S), compared to control samples. Radiation stress probably caused the cell division process to stop at this point without completing cell separation, whereas in control cells, the division was normal.

Furthermore, variations in the cell envelope thickness were frequently observed (see black arrows in Fig. 4). During the highest doses of UVR (15 and 30 min exposure), the different cytoplasmic structures were visualized by electron density variations. In Act20, polyphosphate-like granules (G) were visible, appearing as electron-dense aggregates, scattered throughout the cytoplasm but surrounding the nucleoid (N), with diameters varying between 0.2-0.3 μm (Fig. 4C, 4D); as well as mesosome-like structures (m) formed from projections of the cytoplasmic membrane (Fig. 4D). Also, the interaction between neighboring cells could be observed (see white arrows in Fig. 4D). In NH, the damage caused by UV at the cytoplasmic membrane level was more pronounced. The heterogeneous and disorderly appearance (see grey arrows in Fig. 4G, 4H) could result from loss of membrane integrity, leading to a malfunction of the permeability barrier and inducing cell lysis. Mesosome-like structures (m) (Fig. 4G, 4H) are frequent, and the interaction between bacteria mediated for pili or EPS is a common feature too (see white arrows).

### 3.5. Genome insights in Act20 biotechnological potential

Act20 is a halophile with enormous biotechnological potential, as it encodes haloenzymes and proteins with current applications in the food industry, waste treatment, medicine, cosmetics, biocontrol, pharmacology, paper industry, bioremediation, fuel, and chemical industry (Table 5).

**Table 5.**
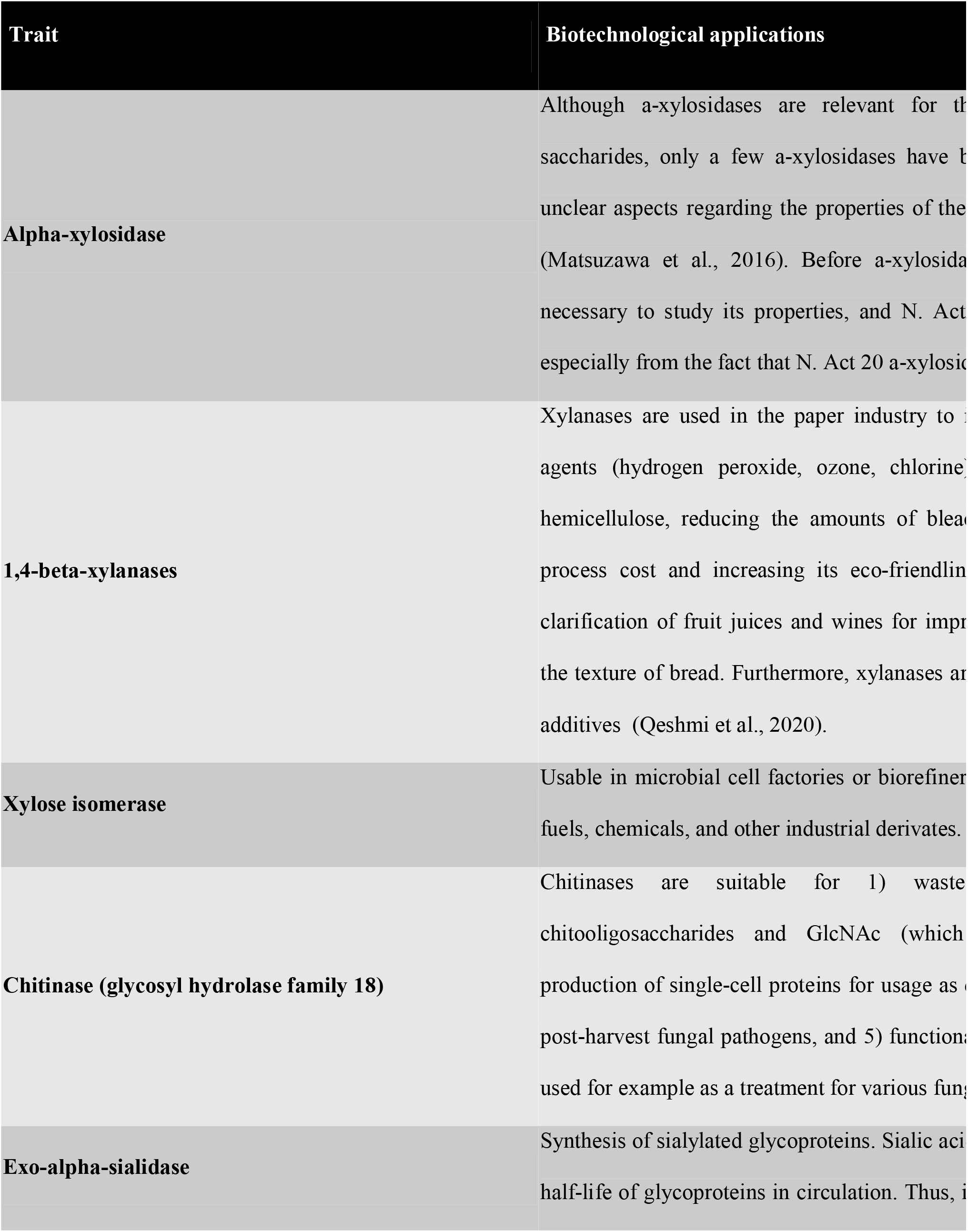
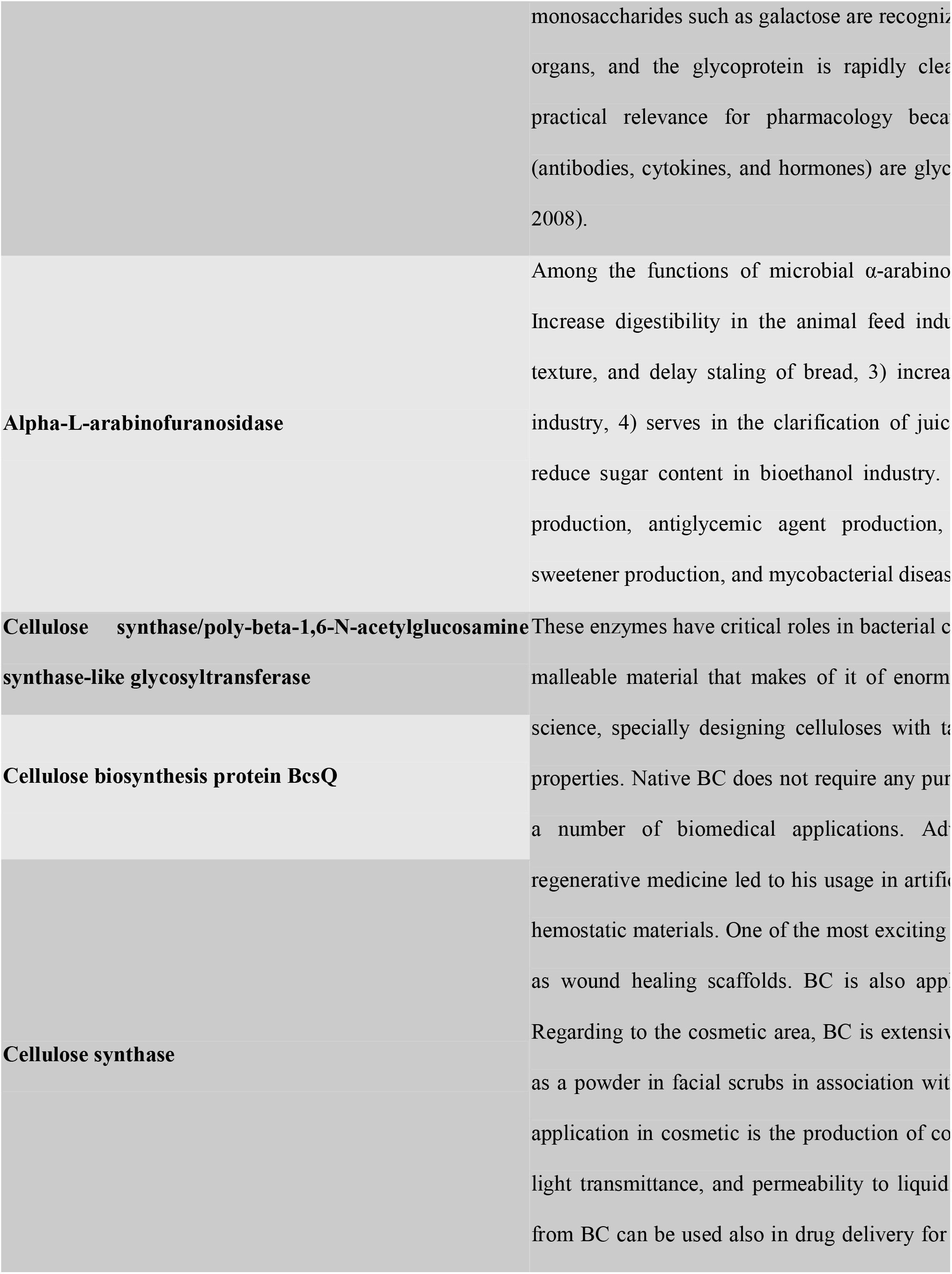
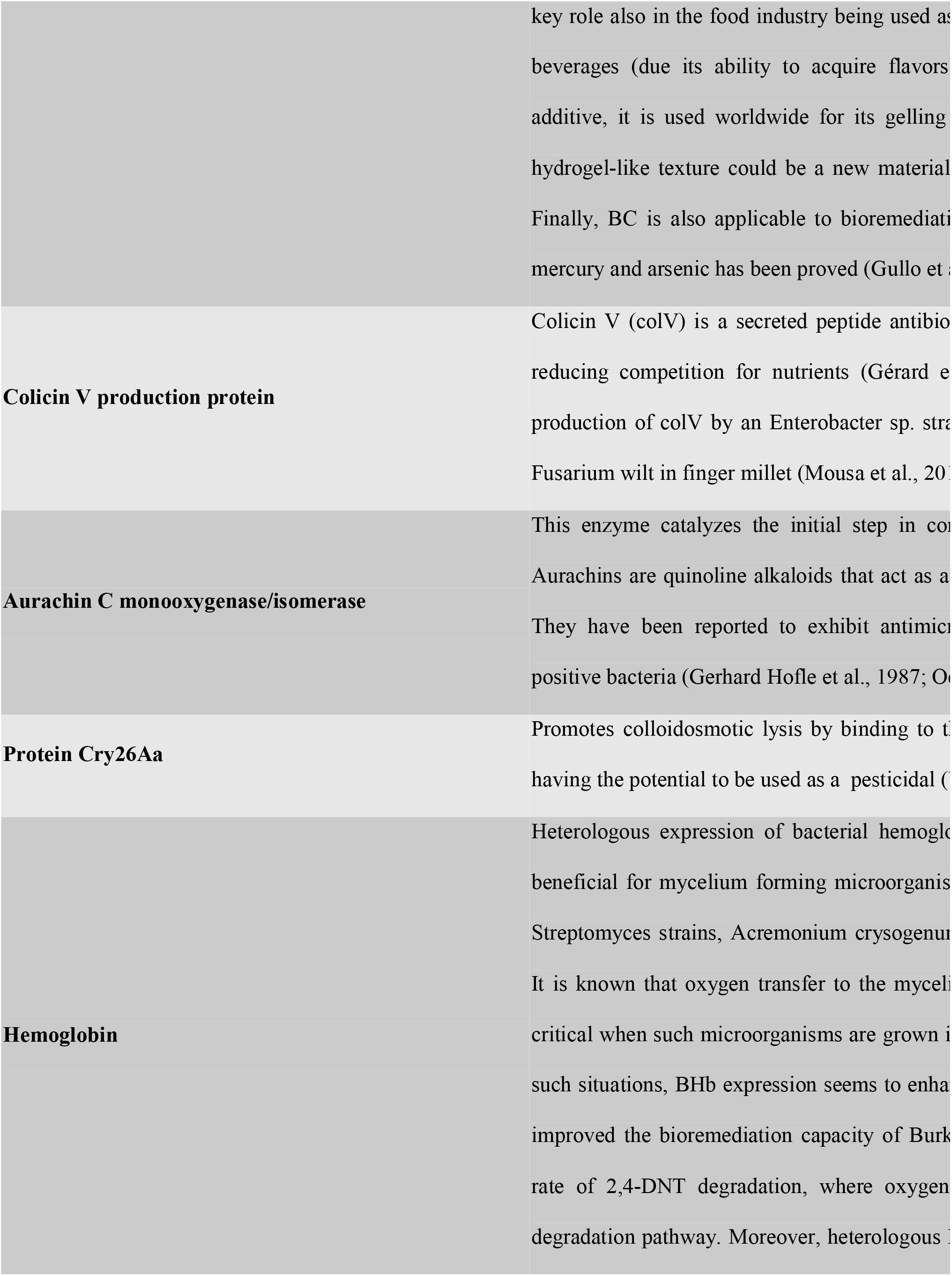
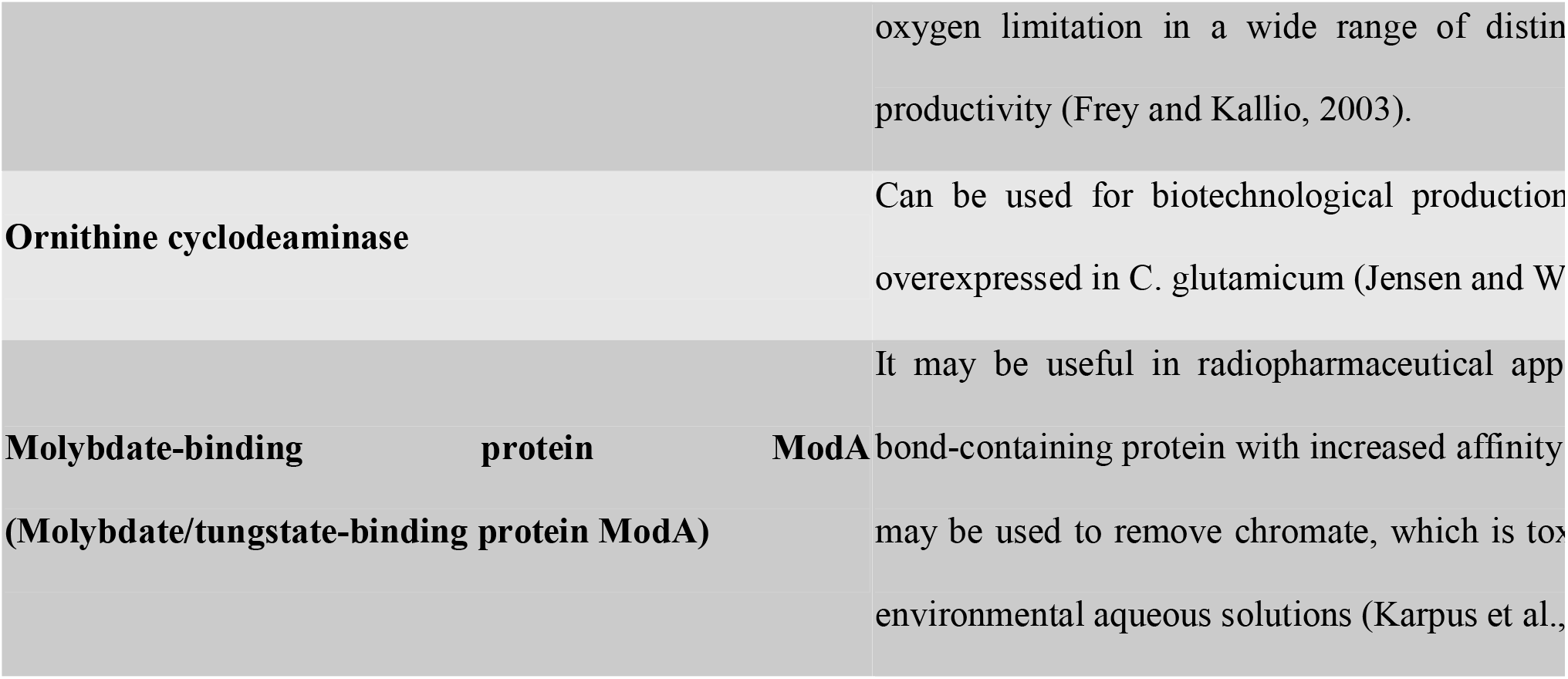
List of Act20’s annotated enzymes and proteins with reported biotechnological potential.

Act20 could degrade wasted vegetal biomass, especially lignocellulose derivatives, as it encodes α-xylosidase, xylanases, xylose isomerases, and an α-arabinofuranosidase. Also has a chitinase, which confers the potential to degrade wasted animal biomass. The genome of Act20 also reports the possibility of producing some exciting compounds of interest; these include bacterial cellulose, a compound with an emerging number of applications such as nanoparticle science and regenerative medicine (Gullo et al., 2018); bacterial hemoglobin, which was previously showed to enhance bioproduction upon low oxygen conditions (Frey and Kallio, 2003), promising antibiotics like colicin and aurachin (Gérard et al., 2005; Gerhard Hofle et al., 1987; Mousa et al., 2016; Oettmeier et al., 1994), and the protein Cry26Aa with high insecticide properties (Wojciechowska et al., 1999). Another relevant enzyme provided by the Act20 genome is the α-sialidase, which has been proven to be useful in synthesizing sialylated glycoproteins (Kim et al., 2011) highly suitable for pharmacology (Varki, 2008).

## 3. DISCUSSION

*Nesterenkonia sp*. Act20 is an actinobacterium isolated from an Andean soil in the Puna region, one of the most extreme environments on the planet, which even NASA has used to test microbes and equipment for spatial explorations (Cabrol et al., 2007; Cockell et al., 2019). Although the polyextremophilic nature of Act20 was preliminary explored before, its genome’s functional characteristics have remained uncovered. In this work, we revealed the genomic basis of the multi-resistance phenotype of strain Act20, especially towards UV radiation, copper and desiccation. Its potential on the production of enzymes and compounds useful for biotechnology was likewise explored.

The ‘extreme’ environmental conditions suffered in its original environmentarid soil at 3,600 m-challenged Act20 to evolve mechanisms to tolerate a wide range of chemical and physical stresses. They include strong fluctuations in daily temperature, hypersalinity, alkaline pH, high levels of UV radiation, a low nutrient availability, desiccation, and high concentrations of heavy metals and metalloids, especially arsenic (Albarracín et al., 2016, 2015; Dib et al., 2008, 2009; Farías et al., 2009; Fernández Zenoff et al., 2006; Ordoñez et al., 2009). *Nesterenkonia* strains were frequently isolated from environmental niches, including other saline locations (Amiri et al., 2016; Yoon et al., 2006). It was demonstrated that the ecology of these diverse habitats defines the genetic differentiation of *Nesterenkonia*. Such genetic differentiation seems to be a key feature in the genome of Act20 as it shares low sequence similarity with other strains of the set of genomes analyzed with ANI. The genomic and physiological particularities suggests Act20 may be a new species adapted to the HAAL environment; experiments heading this way are currently in progress in our lab (Fig. 1).

Act20’s copper and desiccation resistance described before is coincident with several sequences found it its genome potentially involved in such resistance. It is noteworthy the abundance of direct and indirect mechanisms implicated in the copper resistance; some of them were well studied in Gram-negative bacteria and are performed by periplasmic proteins, which are unlike to exist the Gram-positive Act20 which lacks outer membrane and periplasm (Giachino and Waldron, 2020). Likewise, the desiccation resistance relies on the production of a wide diversity of osmoprotectants that creates a hydrostatic force in the cytoplasm; the puzzling fact is that MdoB protein would provoke such hydrostatic pressure only in the periplasm. In both cases, it will be interesting to assess whether Act20 cell wall structure can accommodate a periplasmic-like space where the described gene products can function as proposed for Gram-negative bacteria, periplasmic space was reported for other Gram-positive bacteria (Matias and Beveridge, 2005; Zuber et al., 2006).

A particular cytosolic copper storage protein in the annotation analysis called our attention. Its function and existence are controversial since a widely accepted view is that bacteria have not evolved to use intracelullar copper accumulation due to potential toxicity associated with their metalation (Dennison et al., 2018). Note that this protein was initially discovered in the Gram-negative methane-oxidizing bacterium (methanotroph) *Methylosinus trichosporium* OB3b, which use large amounts of copper to metabolize methane via the membrane-bound (particulate) methane monooxygenase (pMMO) (Vita et al., 2015). Nevertheless, it is not clear the role of this copper storage in Act20 which can otherwise work as an additional osmoprotectant. More research will be needed to confirm the copper bioaccumation ability of Act20, a mechanism useful for the design of bioremediation processes and applicable to several pollutant activities such as industry and mining.

The complete set of genes for ectoine synthetic cluster is present in Act20 genome (Fig. S3), conferring the strain an excellent potential for future biotechnological applications. Ectoine is a water-binding zwitterionic amino acid derivative with numerous biotechnological applications. It is a common component of cosmetic anti-aging and moisturizing creams to improve skin resistance to surfactants in skin cleansing solutions. It also alleviates skin inflammation, being currently recommended to treat moderate atopic dermatitis (Bownik and St pniewska, 2016). Furthermore, the compound is useful in sunscreens as it strongly absorbs ultraviolet (UV) radiation and protects DNA from breaking down in diverse cell types. Ectoine also has applications in medicine: it can inhibit HIV replication and stabilize retroviral vectors for gene therapy (Bownik and St□pniewska, 2016). The alleviation of certain kinds of inflammation (colitis and neutrophilic lung inflammation) and allergic rhinitis were also reported, even preventing the amyloid formation and delaying the onset and progression of Alzheimer’s disease (Bownik and St□pniewska, 2016).

Metabolism of arsenic, a toxic element that can limit or suppress bacterial growth, is also possible for Act20. It has previously been demonstrated that Act20 can tolerate arsenic in the form of As (V) (0-200 mM) (Rasuk et al., 2017). In this work, we reported the presence of an arsenate reductase (not specified), an arsenate-mycothiol transferase, an arsenic transporter (not specified), and seven regulators of the ArsR family (Table S3). We propose that this resistance probably occurs by reducing As (V) to As (III) with a mycothiol-dependent arsenate reductase and subsequent efflux of As (III) from the cell using specific transporters (Ordóñez et al., 2009). Thus, the genome of Act20 joins others already sequenced from HAAL extremophilic prokaryotes (Burguener et al., 2014; Farias et al., 2011; Ordoñez et al., 2015, 2013), being the first of the *Nesterenkonia* genus reported for an environment with a high concentration of arsenic, and the first HAAL genome reporting the reduction of As (V) to As (III) through the mycothiol/thioredoxin redox pathway.

The genome of Act20 also seems to be optimized to cope with low nutrient availability, particularly phosphorous, as genes for a phosphorous-specific two-component system and transporters are well represented (Fig. S4, S5). The two-component system detects low phosphorous availability and communicates the signal to the cell, which expresses phosphate and phosphonate active transporters. Also, two phosphate starvation inducible proteins were reported in the annotation, PhoH, and SphX (Table 4, Table S3), being the latter unique of Act20 and no present in other *Nesterenkonia*. The genomic data also reveals that Act20 could face carbon starvation as its genome codifies active transporters to uptake several kinds of peptides, nucleotides, amino acids, and carbohydrates from the environment (Table S3, Fig. S4).

*Nesterenkonia* sp. Act20 genome is the first one of this genus reported from the highest UV irradiation environment on Earth: Puna-High Andes region; in accordance, we found genetic traces for complex UV resistance mechanisms comprehensively called as UV-resistome. The comparison of UV-genomic determinants of Act20 with other *Nesterenkonia* genomes indicated indeed a more sophisticated UV-resistome for the Socompa strain. It is then evident that environmental irradiation has a notable impact on the genome of Act20, which has a higher quantity and diversity of genes dedicated to UV resistance than other strains less irradiated. We also observed a similar pattern for other *Nesterenkonia* from high altitude or expected high irradiated environments (Fig. 2, Table S7). Interestingly, solar irradiation intensity selecting UV-resistome gene abundance and diversity in aquatic microbiomes was also evidenced by our group using a worldwide metagenomic analysis (Alonso-Reyes et al., 2020).

Among the subsystem of UV evasion/shielding is worth to note the genes involved in the production of gas vesicles (gvpA, K, O and F). Certainly, the expression of gas vesicles (Damerval et al., 1991; Englert et al., 1992; Pfeifer, 2012) along with flagella, allow microbes to move up in the water column toward sunlight. However, in many cases, these vesicles are also present in soil prokaryotes, even from high UV radiation environments, thus suggesting new roles in radiation protection (Oren, 2012). It has been speculated that they could change the cell position concerning the angle of the light’s incidence, changing its impact on the cell (Bolhuis et al., 2006; Oren, 2012). Additionally, our current works on comparative proteomics reveal an increase in vesicle proteins’ expression in response to the UV increase (Zannier et al., in preparation). We also include flagella, swarming motility proteins and the pili, whose ability to promote bacterial aggregation or biofilm adhesion may protect the cells from the UV exposure (Burdman et al., 2011; Ojanen-Reuhs et al., 1997).

Act20 genome codes resistance genes to other extreme factors: low temperatures, low atmospheric O_2_ pressure, heavy metal and other toxic compounds stress. Many of these hard environmental factors added to the previously mentioned and geophysical characteristics of the sampling site resembles those present in Early’s Earth atmosphere that witnessed the evolution of ancient microorganisms (Albarracín et al., 2015; Cabrol et al., 2007) (Fig. 5) (Cockell et al., 2000; Forni et al., 2015; Hecht et al., 2009; Karunatillake et al., 2007; Sforna et al., 2014; Wadsworth and Cockell, 2017; Yen et al., 2006) (https://mars.nasa.gov/all-about-mars/facts/). Thus, Act20 is an exciting model to study the mechanisms by which the extremophiles could have successfully faced the adverse conditions of the Earth’s history, having clear implications on astrobiological projects (Hiscox and Thomas, 1995; Merino et al., 2019; Slotnick, 2000).

**Figure 5.**
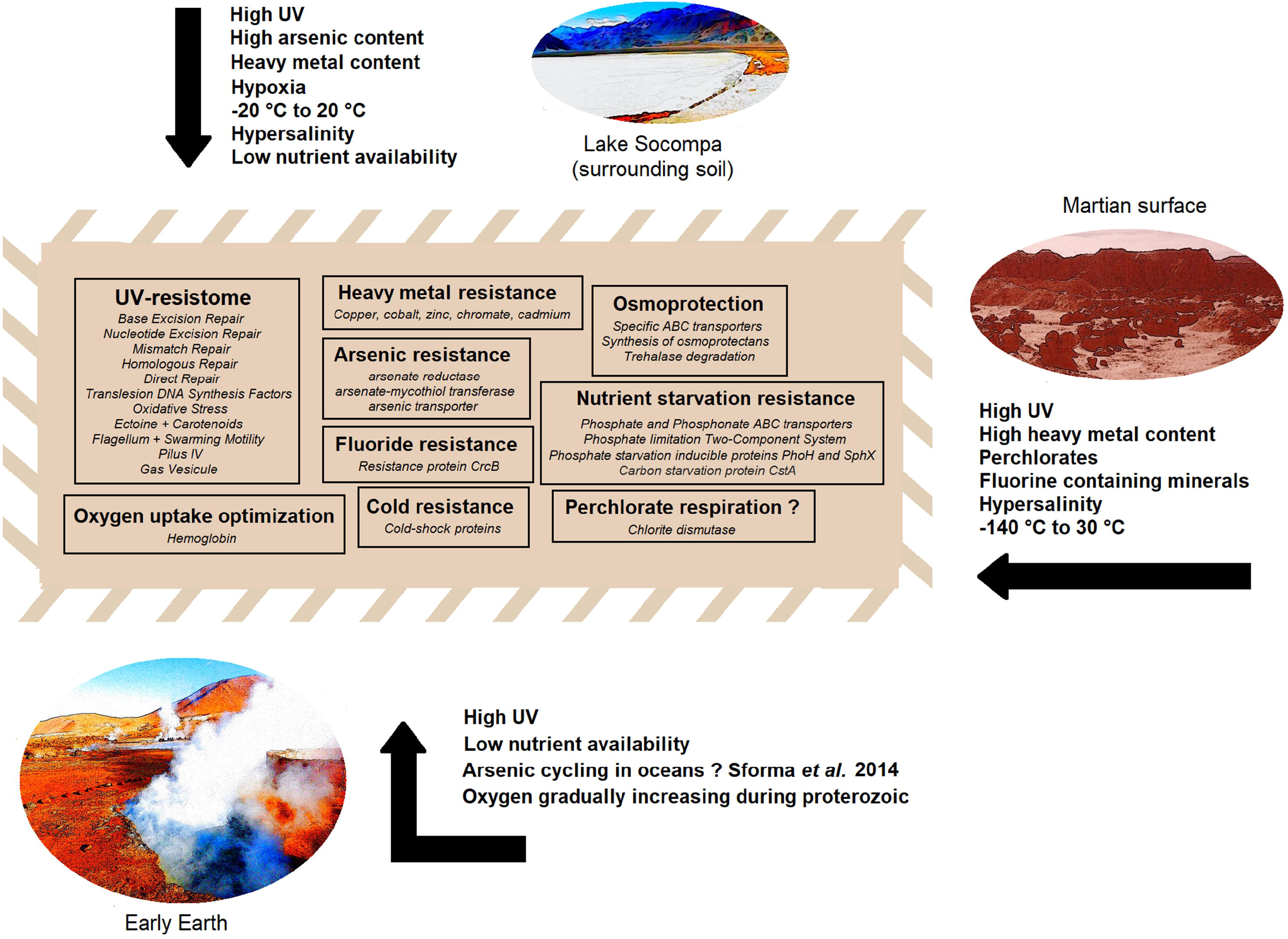
Comparison between the extreme conditions (relevant to Act20) of the Lake Socompa, Mars surface, and the early Earth environments. The light brown box represents the Act20 genome and its potential for the different resistances as described inside.

Act20 could also have a great biotechnological potential for producing and metabolizing compounds and enzymes of interest (Table 5). Bacterial cellulose, hemoglobin, antibiotics, and a potential insecticide are among the Act20 biosynthetic products. On the other hand, several genes for enzymes capable of degrade vegetal biomass were also detected. This feature is of importance for the biodegradation of lignocellulosic biomass of agro-industrial wastes which are produced in large amounts through agricultural and forestry practices, including the paper-and-pulp and timber industries. On the other hand, animal biomass that contains high proportions of shellfish, such as shrimp, crab, and krill, are suitable to be processed through chitinases, an enzyme that is also present in Act20. The seafood processing industry has raised serious concerns regarding disposal issues because of this waste’s low biodegradation rate, which could be solved through an enzymatic approach.

The enzymes mentioned above could be used in industrial processes that reproduce the microbe’s original natural habitat’s extreme conditions. Biocatalysts isolated by these extremophiles are termed extremozymes and possess special salt allowance, thermostability, and cold adaptivity. Extremozymes are very resistant to severe conditions owing to their great versatility. As such, they represent new prospects for biocatalysis and biotransformations and the development of the economy and unique line of research through their application (Dumorné et al., 2017). Here, we also report the genetic potential of Act20 to provide poly-extremozymes, which could combine resistance to cold (−2 ° C - 20 ° C), high solar radiation, salinity (at least 1 M salt), and high pH (>8). Enzymes of this halophilic microbe could provide great opportunities, particularly for food, bioremediation, and pharmacy industries.

## CONCLUSION

In this work, we have confirmed by lab assays the multi-resistance phenotype of *Nesterenkonia* sp. Act20, a poly-extremophile originally isolated from Puna arid soil surrounding Lake Socompa, in the Puna region, exposed to the highest irradiated environment on Earth. Accordingly, its genome codes for a plethora of genes that help counteract the ecological pressure of the hostile conditions face by the microbe in its original environment: i.e. arsenic, nutrient limiting conditions, osmotic stress, UV radiation, low temperatures, low atmospheric O2 pressure, heavy metal and other toxic elements stress.

As a novel extremophile, Act20 has the potential to produce compounds (extremolytes and extremoenzymes) of interest with application in industrial processes such as an insecticidal protein, bacterial cellulose, ectoine, colicin V, arauchins, chitinases and cellulases. In this way, Act20 becomes an exciting candidate for additional studies of transcriptomics, proteomics (currently in progress), metabolomics, as well as the expression and testing of biotech-competent enzymes. The herein report shed light on the microbial adaptation to high-altitude environments, its possible extrapolation for studying other extreme environments of relevance, and its application to industrial and biotechnological processes.

## 4. ACKNOWLEDGEMENTS

The authors acknowledge the generous financial support by PIUNT G603 and PIP CONICET 0519 projects. VHA, MPV, and MEF are staff researchers from the National Research Council (CONICET) in Argentina. DA, MSG, NNA are the recipients of doctoral fellowships from CONICET. Act20 genome sequencing project was performed in INDEAR-CONICET, Argentina. Electron micrographs used in this study were taken at the Center for Electron Microscopy (CIME) belonging to UNT and CCT, CONICET, Tucuman. This manuscript has been released as a Pre-Print at bioRxiv.

## Notes

### Competing Interest Statement

The authors have declared no competing interest.

