## Supplementary figures and images for "Genomic Insights of an Andean Multi-resistant Soil Actinobacterium of Biotechnological Interest"

### Fig. S1

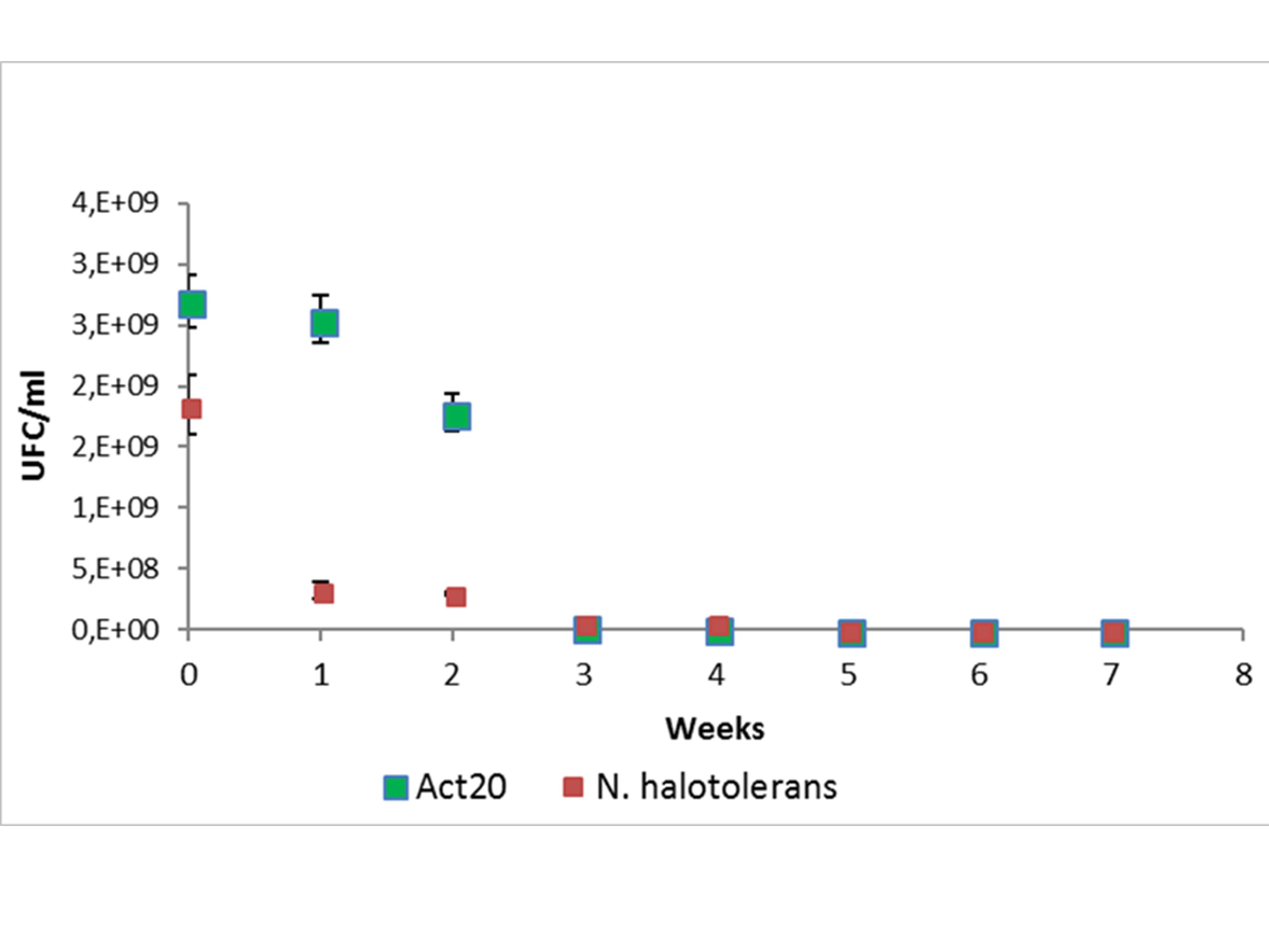

### Fig. S2

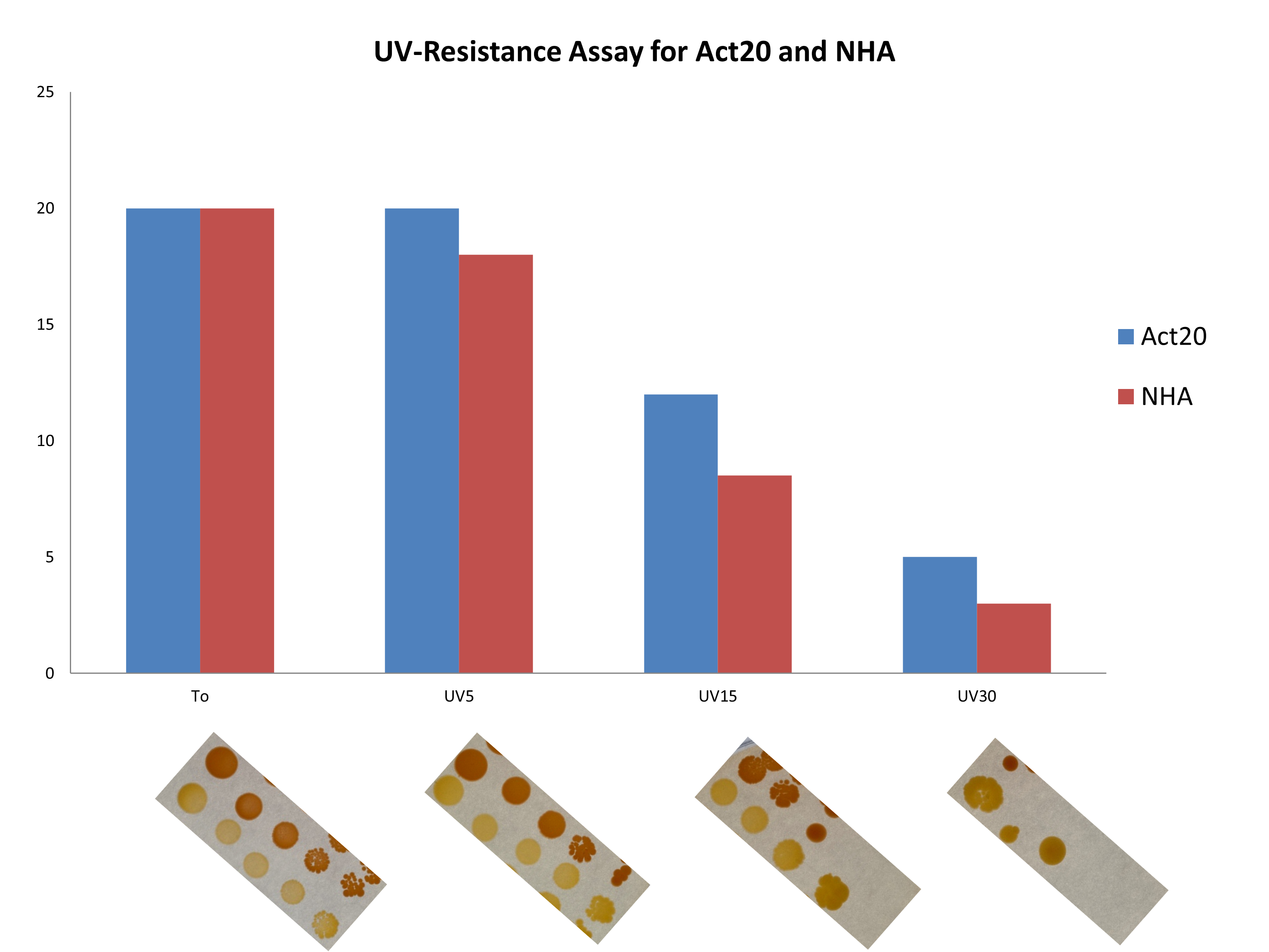

### Fig. S3

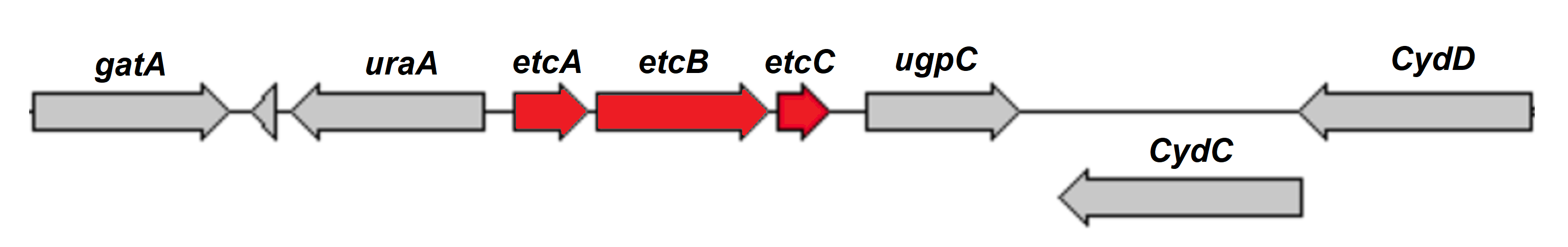

### Fig. S4

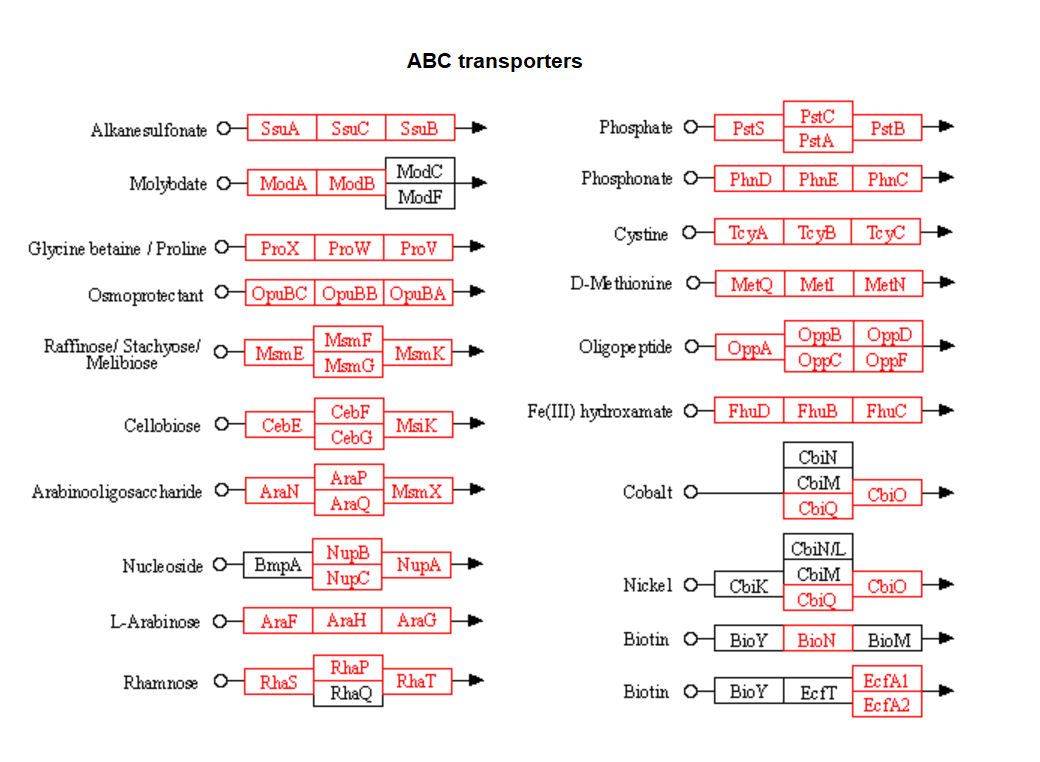

### Fig. S5

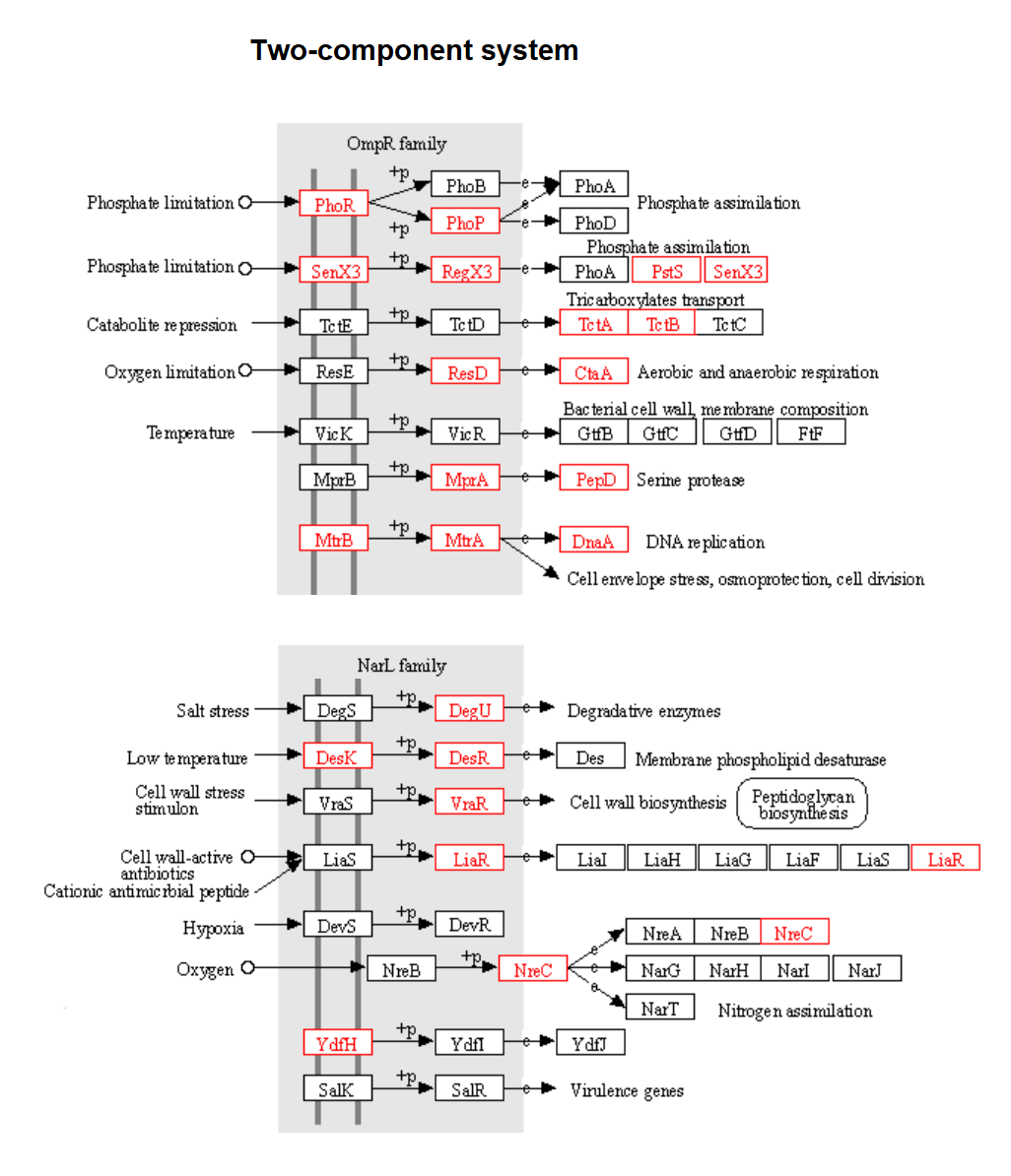
